# Fermentative yeast diversity at the northern range limit of their oak tree hosts

**DOI:** 10.1101/2025.01.30.635702

**Authors:** Javier Pinto, Chloé Haberkorn, Markus Franzén, Ayco J. M. Tack, Rike Stelkens

## Abstract

Fermentative yeasts play important roles in both ecological and industrial processes but their distribution and abundance in natural environments is not well understood. We investigated the diversity of yeast species associated with oak trees (*Quercus spp*.) in the northern range limit of oak in Sweden, and identified climatic and ecological conditions governing their distribution. We isolated yeasts from oak bark collected from 28 forests and identified them to the species level using metabarcoding. Most communities were dominated by species in the Saccharomycetaceae family, especially by species of *Saccharomyces, Kluyveromyces*, and *Pichia*. Each of these genera showed a distinct latitudinal and longitudinal distribution, and both temperature and precipitation metrics predicted significant variation in their abundance. Consistent with this, laboratory assays revealed significant effects of temperature on the growth of strains collected from different longitudes and latitudes. We further found that older trees harbour more diverse and more balanced microbial communities with more evenly distributed species abundances, and that communities across trees were more similar when sharing a common dominant genus. This work provides a baseline for future studies on the impact of climate change on the microbial biodiversity of temperate forests in northern latitudes, and contributes to a growing collection of fermentative yeasts isolated from the wild with potential for biotechnological applications.

## Introduction

Fermentative yeasts are abundant in temperate forests where they play a crucial role in ecological processes (Mozzachiodi et al., 2022). They recycle organic matter, mediate nitrogen and carbon cycles, and serve as a food source for insects (Botha, 2011). They also impact microbial community composition, e.g. by inhibiting the growth of competing microorganisms by producing ethanol (Viljoen, 2006). Fermentative yeasts are also known for their high stress tolerance and their efficient use of monosaccharides and nitrogen (Treseder & Lennon, 2015; Romero-Olivares et al., 2021), making them well-suited for industrial applications, particularly for the production of alcoholic beverages and biofuels. Recent studies suggest that the metabolic diversity of natural yeasts, their variation in niche breadth and stress tolerance, is largely shaped by genetic factors (Opulente et al., 2024). A recent global distribution analysis showed that species richness is highest in mixed, montane forests in temperate climates, and is best predicted by microhabitat, vegetation type, and topography (David et al., 2024). Species ranges are strongly influenced by overlaps with other yeast species, indicating that niche partitioning plays an important role in their biogeography. The distribution of budding yeast in the genus *Saccharomyces* is known to be associated with environmental temperature (David et al., 2024; Mozzachiodi et al., 2022; Sweeney et al., 2004).

Oak trees (*Quercus spp*.) have been described as a frequent habitat for fermentative yeast, in different parts of the world (Mozzachiodi et al., 2022; Robinson et al., 2016; Sampaio & Gonçalves, 2008; Sniegowski et al., 2002). Oak trees, especially their bark and sap, provide natural sugars (Ferreira et al., 2018) and can accumulate decaying plant matter and moisture, which creates suitable growth environments for yeast. Once the polysaccharides in the bark are broken down into smaller sugars, e.g. by filamentous fungi (Battaglia et al., 2011; de Vries & Visser, 2001), they are available as a carbon source for fermentative yeasts. Forests in the north of Europe represent the northern range limit of oak, which is marked by long, cold winters and warm (but not hot) summers, a shorter growing season, and more coniferous vegetation. Historically, oak forests were widespread in southern Scandinavia. Over the past centuries, a combination of biogeographical shifts and human activities have significantly reduced the volume of oak forests (Löf et al., 2016). Tree health has also substantially deteriorated, especially in *Quercus robur*, which has suffered from crown defoliation and a general decline in recent decades (Drobyshev et al., 2007). Today, the remaining oak forests are concentrated along the coastlines in southern Sweden in the temperate and hemiboreal climate zones, delimited in the north by the subarctic climate of the boreal zone (Drobyshev et al., 2008). Although remaining oak forests in Sweden are likely important reservoirs of microbial diversity, the current lack of microbial data has precluded a systematic investigation of this ecosystem. Whether fermentative yeasts are part of the oak microbiome also in the northernmost range of the *Quercus* distribution, is so far unknown.

Our overarching aim was to investigate the diversity of fermentative yeasts found in the northern range limit of oak, and identify climatic and environmental drivers of their distribution. For this, we isolated yeasts from oak bark samples collected across southern Sweden, specifically from the nemoral and hemiboreal climate zones (Jonsson et al., 2016). Our sampling area covered 583 km from south to north, along latitudes between 55°52’33’’N and 60°76’78’’N (from southern Sweden to as far north as the Swedish oak tree line allowed), and 379 km from west to east, along longitudes between 12°48’38’’E and 18°63’15’’E (from the North sea to the Baltic sea). We used enrichment protocols designed for fermentative yeast in the genera *Saccharomyces, Komagataella, Lachancea*, and *Candida* (Cubillos et al., 2019; Sampaio & Gonçalves, 2008; Sniegowski et al., 2002; Villarreal et al., 2022). For biodiversity and abundance assessment, we used metabarcode sequencing of the internal transcribed spacer (ITS) region, allowing identification down to the species level (Alsammar et al., 2019; Větrovský et al., 2019). Specifically, we asked 1) if microbial communities differ in their composition and diversity, 2) if there are spatial patterns in microbial community structure along latitudinal and longitudinal gradients, reflecting variation in temperature and precipitation, 3) if oak tree hosts characteristics (e.g. tree age) predict yeast diversity, and 4) if the growth of the yeast strains we isolated is affected by temperature using laboratory assays.

## Materials and Methods

### Yeast sampling from oak tree bark

Eight oak trees were sampled in each of 28 stands, at 23 different locations across the south of Sweden, in both nemoral and hemiboreal vegetation zones (**Figure 1**). Trees included 67 *Q. robur*, nine *Q. petraea*, and five oak trees (*Q. sp*) with unidentified species status. Sampling period was during the mid and late summer, from July to September 2023. Approximately 15g of bark (ca. 4 × 1× 0.5 cm), including both the outer and inner bark layers, was collected from each tree using an ethanol-sterilized knife. Where single pieces of this size were not obtainable, multiple smaller bark pieces were collected. Samples were collected using nitrile gloves and stored individually in sterile plastic bags. All sampling tools were sterilized with 90% ethanol before and after sampling. Samples were transported in a cooled container and stored refrigerated until processing.

**Figure 1.**
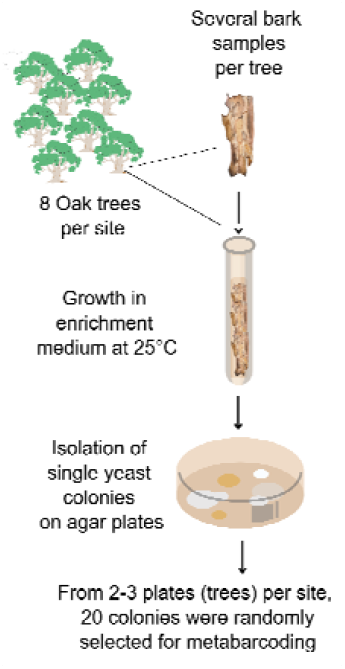
Schematic of the sampling, enrichment, and isolation of yeast from oak trees.

Bark samples from each tree were grown in 25 mL of liquid enrichment medium (a modified version of the medium described by Sampaio & Gonçalves, 2008), consisting of YNB (yeast nitrogen base) supplemented with 1% (wt/vol) raffinose and 4% (vol/vol) ethanol. Instead of using 8% ethanol, we reduced ethanol to 4% to increase our chances of uncovering more biodiversity, including groups of yeast that are not able to tolerate high alcohol concentrations. We inoculated for 15 days in 50 ml Falcon tubes at room temperature. From each tube, 100 μL of liquid was then sampled and diluted 4X in sterilised water. Fifty microliters of diluted sample were then plated onto solid YMA media (1% glucose, 0.5% peptone, 0.3% yeast extract, 0.3% malt extract, 2% agar) with 4% ethanol using glass beads, and grown for five days at 30°C. To confirm the presence of yeast, a PCR of the ITS region was performed for three colonies per stand (**Figure S1**). Two to three individual trees per stand with growth of putative yeasts were then randomly selected for DNA extraction (total number of trees n = 81).

### DNA Extraction

From each enrichment plate (one per tree) we randomly selected ∼20 colonies, scraped the colonies with a sterile pipette tip, and added them to 250 μL of 1X PBS. After vortexing, samples were spun down at 3000 rpm for three minutes and the supernatant was discarded. Fifty microliters of zymolyase solution (1 mg/mL zymolyase, 1 M sorbitol, pH 7) were added to the pellet and mixed by pipetting. Cells were then incubated at 37°C for 2h before adding 1 mL of sterilised water. Cells were then spun down at 13 000 rpm for 30 seconds to discard the supernatant and resuspended in 250 μL PBS. After vortexing, 20 μL of proteinase K solution and 290 μL of AL buffer were added to each sample, and mixed thoroughly by pipetting. Samples were then incubated at 60°C for 20 minutes and cellular debris was removed by centrifugation at 3,000 rpm for 3 minutes. A KingFisher 96 deep-well plate was prepared following the OMEGA Bio-Tek protocol to perform the DNA extraction. PCR1 was performed to enrich the samples for putatively present yeast DNA, by amplifying the ITS region using primers for ITS4 and ITS5 (**Figure 2**, primer sequences in **Table S1**) (White et al., 1990; Alsammar et al., 2019). Amplification was carried out using the Takara Taq polymerase kit. PCR1 amplicon concentrations were assessed using Qubit dsDNA HS Assay Kits. All concentrations were normalised to 2ng/μL by diluting samples in MilliQ water, up to a volume of 50 μL per sample.

**Figure 2.**
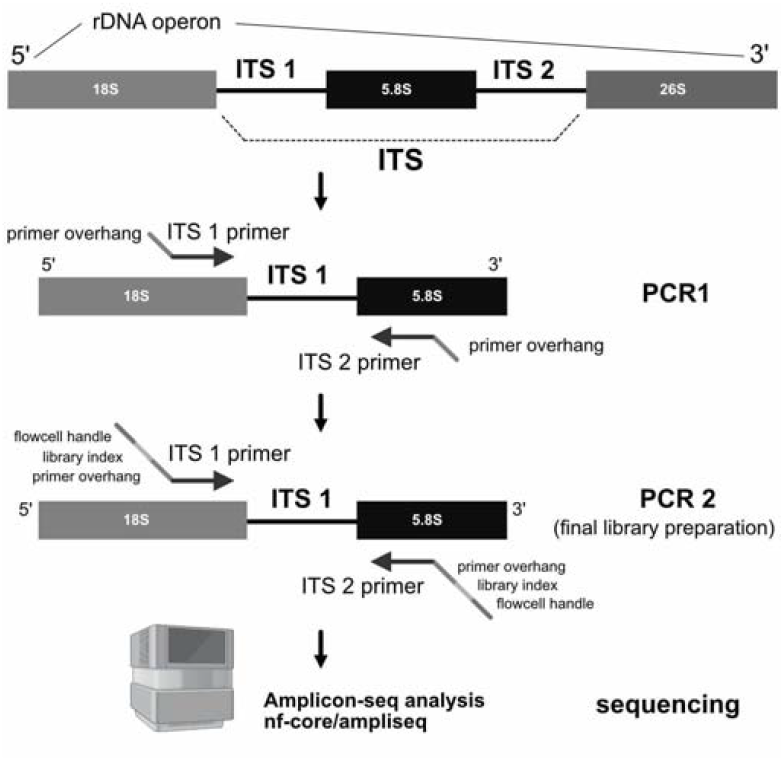
Schematic of ITS amplification using PCR1 with primer overhangs (**Table S1**), followed by indexing in PCR2 for Illumina library preparation and sequencing.

### Sequencing

PCR1 amplicons with normalised concentrations were cleaned up using MagSI-DNA NGS PREP Plus magnetic beads at the National Genomics Infrastructure (NGI, Solna, Sweden), followed by an indexing PCR2 with Adaptamera indices and a second cleanup using magnetic beads. Quality control was performed on the final library before sequencing on the NovaSeq 6000 platform. Raw reads were deposited on NCBI’s Sequence Read Archive (SRA) database under BioProject PRJNA1114957.

### Read Processing

Reads were processed to identify yeast species using the nf-core/ampliseq pipeline (Straub et al., 2020) with default parameters and option ‘illumina_pe_its’. Amplicon Sequence Variants (ASVs), i.e. unique DNA sequences obtained from amplicon sequencing, were inferred and annotated with DADA2 against the UNITE general FASTA release for Fungi v9.0 (Abarenkov et al., 2024). ASVs were kept when their sequence similarity was above a common 97% confidence threshold (Kauserud, 2023). All the scripts used for data processing, analyses and figures are openly available on GitHub: https://github.com/chaberko-lbbe/yeast-oaktree.

### Environmental predictors

Longitudes and latitudes were extracted from GPS coordinates for each sampling location. Using these coordinates, temperature and precipitation data were extracted from WorldClim version 2.1 (Fick & Hijmans, 2017) with QGIS software (Geographic Information System). Average monthly values were computed, and used to extract annual mean, maxima, and minima for both temperature and precipitation at each location. Because means followed the same trends as minima, mean data were not used. Mean annual temperatures ranged between -8°C and 25°C, and precipitation ranged between 13.3mm and 136.7mm, averaged across monthly mean minima and maxima.

### Individual host tree metrics

Tree height was measured using a Suunto PM-5/1520 clinometer. Diameter at 1.3 m breast height (DBH) was assessed using Haglöf Mantax Blue Klave callipers. Bark depth was measured as the mean depth of bark crevices in the four cardinal directions (North, South, East, and West) using a metal ruler, following the methodology of Johansson et al. (2009). Tree age was determined by counting the total number of annual growth rings in each core sample obtained at breast height, following standard dendrochronological procedures and cross-validation (Holmes, 1983). This provided a minimum age estimate, as core sampling at 1.3 m height does not account for the time required for trees to reach breast height. Annual radial growth rates were quantified through tree-ring widths (TRW) measurements using a digital LINTAB positioning table connected to an Olympus stereomicroscope. These measurements were recorded to the nearest 0.01 mm using TSAPWin Scientific software (Rinn, 2003). Each annual ring width represents the radial stem increment for that particular year, providing a high-resolution time series of growth patterns. Growth rates were calculated as the annual increment in ring width, with the mean annual growth rate determined by averaging these measurements across the entire core sample. To account for age-related growth trends and ensure data quality, all TRW series underwent standardization and cross-dating procedures using COFECHA software (Johnson & Abrams, 2009).

### Growth-at-temperature assays

To test whether temperature has an effect on the growth of the strains we isolated from the wild, we assessed the genus affiliation of three colonies per site by Sanger sequencing. Since the strains were collected blindly from the remaining colonies on the plates after amplicon sequencing, some did not match the dominant species identified for each location by amplicon sequencing. Thus, for the growth assays, we only retained strains identified by Sanger sequencing as *Saccharomyces, Kluyveromyces* and *Pichia*, from a total of 18 locations. Sanger sequencing confirmed the presence of *Pichia* in only one sample (Halmstad 1), which precluded us from testing for effects of latitude and longitude on growth-at-temperature for this genus.

We measured the maximum biomass of each strain as a proxy for growth at three different temperatures: 5°C (below 4°C growth is usually no longer observed in yeast; Salvadó Z. et al., 2011), 16°C, which is close to the mean temperature during Swedish summer; and 35°C, which is close to the highest temperature ever recorded in Sweden (SMHI, 2023). Growth assays were performed on a BioTek Epoch 2 microplate spectrophotometer (Agilent, USA) using maximum biomass, measured as optical density (OD_600nm_). We standardised all inoculates to an initial OD_600nm_ of 0.1 (approximately 10^6^ cells) in 200 μL of YPD media (20g/L peptone, 10g/L yeast extract, 2% glucose). Before final growth measurements were taken, each strain was grown for 24 hours (for measurements at 16°C and 35°C) and for 120 hours (for measurements at 5°C) in 96-well plates. We used 10 technical OD replicates per strain. Biomass was blank-corrected using negative controls (6 wells with media but no yeast).

### Statistical analysis

Abundance and taxonomy tables based on the ASV read data were used to compute alpha diversity indices: ASV richness (the number of different ASVs), Shannon (accounting for both ASV richness and their relative abundance; Shannon & Weaver, 1963), and Evenness indices (how evenly distributed ASV abundance is among species within a community; Wilsey & Potvin, 2000) with the package R/vegan v2.5-7 (Oksanen et al., 2020). A non-metric multidimensional scaling (NMDS) ordination was computed based on ASV richness using the vegan function metaMDS (distance “bray”, 2 dimensions, try max 1000). To evaluate whether pairs of species co-occur more or less frequently than would be expected by chance across a set of locations, R/cooccur v1.3 was used (Veech, 2013; Griffith et al., 2016). To test if ASV abundance across the three dominant genera was predicted by geospatial and environmental metrics, we ran linear models using the function lm in R/stats v4.2.3. Pairwise comparisons of slopes between genera were made within each spatial gradient, using the function emtrends (t-test) in the package R/emmeans v1.8.1 (Lenth, 2022). To assess differences in community composition across tree species and insularity, a Bray-Curtis distance matrix was computed based on genus-level abundance data using the vegdist function from the package R/vegan (method “bray”). Permutational multivariate analysis of variance (PERMANOVA) was then conducted with the adonis2 function (R/vegan), using either tree species or insularity as the explanatory variable (999 permutations). Growth-at-temperature assay data were analysed by running linear models against longitude and latitude for each genus at each temperature.

## Results

A total of 149 ASVs were detected, of which 69 were attributed to the genus *Saccharomyces*. Up to 26 different ASVs were detected per tree with on average 166 620 ASV reads per tree. No ASVs were detected in 3 out of 81 trees (one tree each in Blå Jungfrun, Björnstorp and Vårgårda-1).

### 1. Microbial communities across the sampling area differ in composition and diversity

First, we set out to test if microbial communities across the sampling area differ in their composition and diversity. We identified a total of 14 genera of yeasts (**Figure 3A**) belonging to five families across all samples (Debaryomycetaceae, Pichiaceae, Saccharomycetaceae, Saccharomycetales fam. incertae sedis, and Saccharomycodaceae). Computing the sum of ASV reads detected across all samples for each family revealed that species belonging to Saccharomycetaceae were the most frequently detected (71.88%), followed by Pichiaceae (24.94%) (**Figure S2**). Among genera, *Saccharomyces* was the most common (34.06%), followed closely by two other genera of the Saccharomycetaceae family: *Kluyveromyces* (33.91%) and *Pichia* (22.39%) (**Figure 3A**). These three genera represented up to 90.36% of the total ASV reads, while each of the eleven remaining genera represented less than 3% of ASV reads.

**Figure 3.**
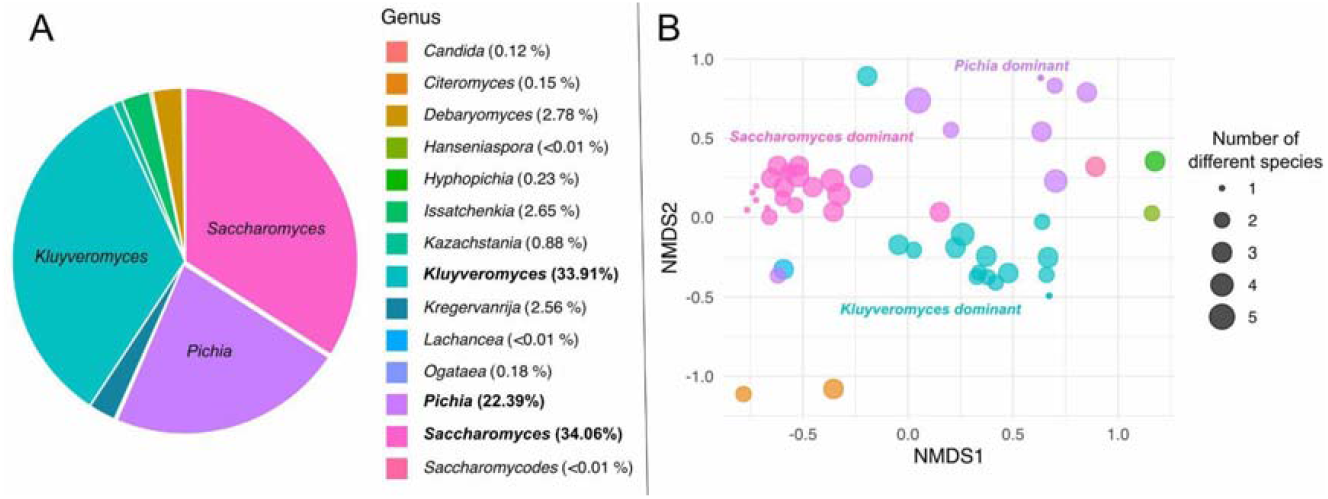
Overview of the 14 yeast genera detected on oak trees in Sweden. **A)** Percentage of ASVs mapping to each genus across all samples. **B)** Projection using non-metric multidimensional scaling (NMDS), where each circle represents a single tree. Ordination is based on ASVs richness with colours indicating the dominant genus. Outlier samples LANY-1, TAN-1 and VASV-1 were removed for display purposes.

Samples were plotted using non-metric multidimensional scaling (NMDS) and coloured according to the most frequent genus (based on the number of ASV reads) within each sample (**Figure 3B**). Three main clusters were revealed, which grouped together the samples that were dominated by species of either *Saccharomyces, Kluyveromyces* or *Pichia* (with a larger spread in the *Pichia* cluster). The proximity of points within each cluster suggests that microbial communities are more similar when sharing a common dominant genus. While there was no significant difference between the three clusters in species richness (median of 2.5 species), the number of yeast species per tree was slightly higher when the dominant genus was *Pichia* (**Figure S3**).

Up to 18 species were found across the 14 genera (**Table 1**). Within the genus *Saccharomyces*, only *S. paradoxus* was detected. In the genus *Pichia*, we identified *P. mandshurica, P. membranifaciens*, and *Pichia sp*.. In the genus *Kluyveromyces*, we found *K. lactis* and *K. dobzhanskii*.

**Table 1.**
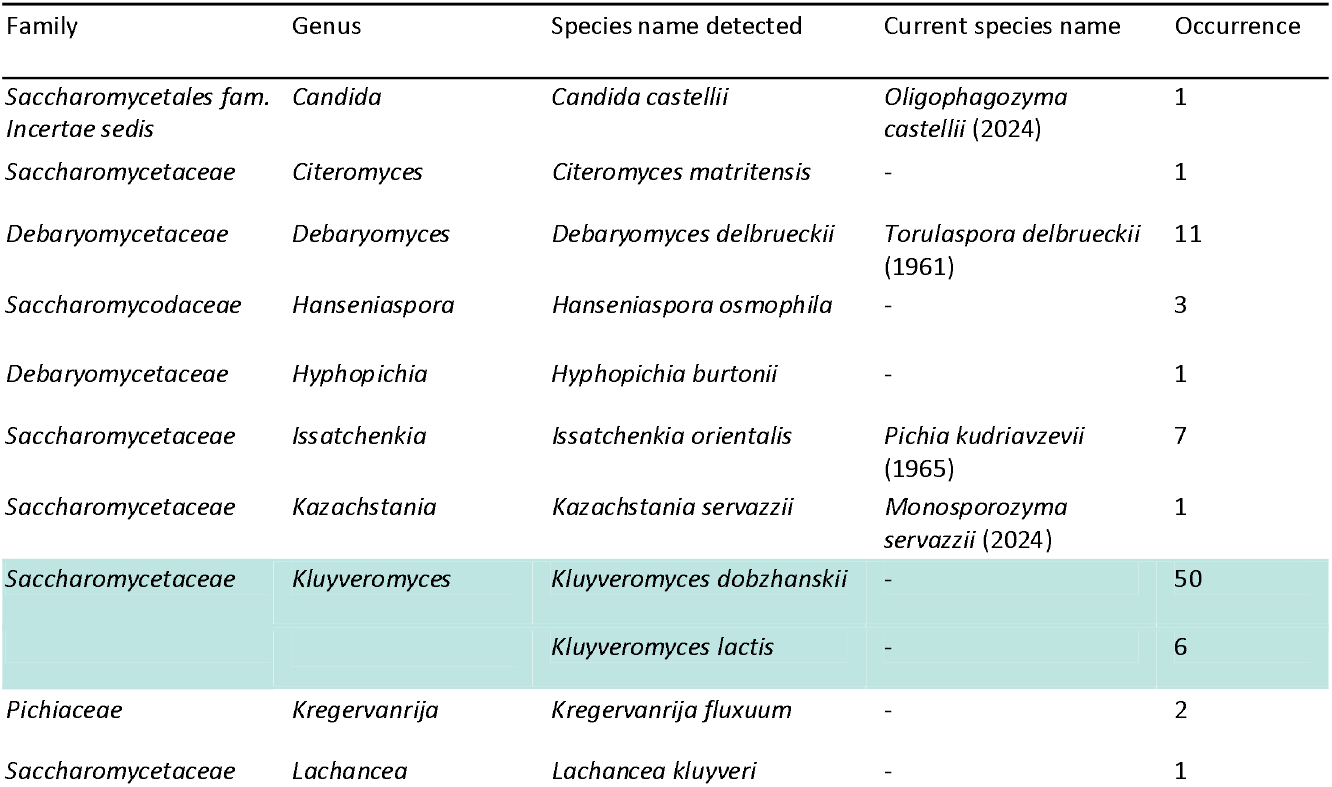
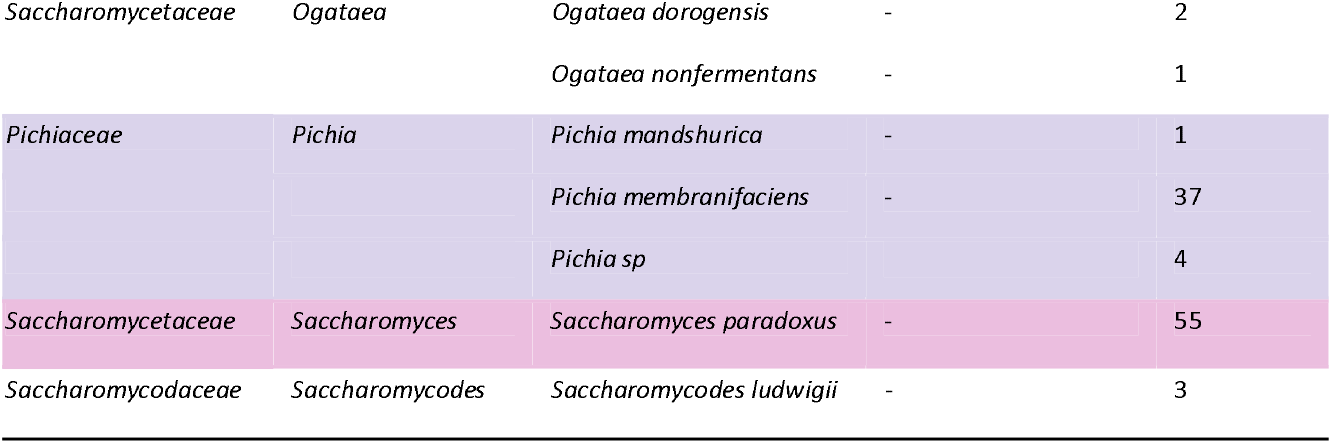
Species detection across all genera. Occurrence represents the number of trees (out of 81) where species were detected with at least one ASV. Species names were assessed using Fungi v9.0, and current names were extracted from MycoBank (Robert et al., 2013).

Co-occurrences of the 18 yeast species detected were visualised using a chord diagram (**Figure 4**). The thickness of the connecting lines between species reflects the frequency with which they co-occur across the sampled locations (between zero and 34 times; **Figure S4**). The species that co-occurred most often were *Saccharomyces paradoxus, Kluyveromyces dobzhanskii* and *Pichia membranifaciens*. However, when comparing the observed frequency of co-occurrence to what would be expected under a random null model (binomial test), only a single significant positive co-occurrence was detected (p-value ‘greater than expected’ < 0.05) between *Debaryomyces delbrueckii* and *Saccharomyces paradoxus*, suggesting that these species are more likely to co-occur, possibly sharing compatible ecological requirements or forming a symbiotic relationship. On the other hand, *Pichia sp*. and *Saccharomyces paradoxus*, as well as *Debaryomyces delbrueckii* and *Kluyveromyces dobzhanskii*, exhibited significant negative co-occurrence (p-value smaller than expected’ < 0.05) suggesting that these species are less likely to be found in the same environment due distinct habitat preferences or niche competition.

**Figure 4.**
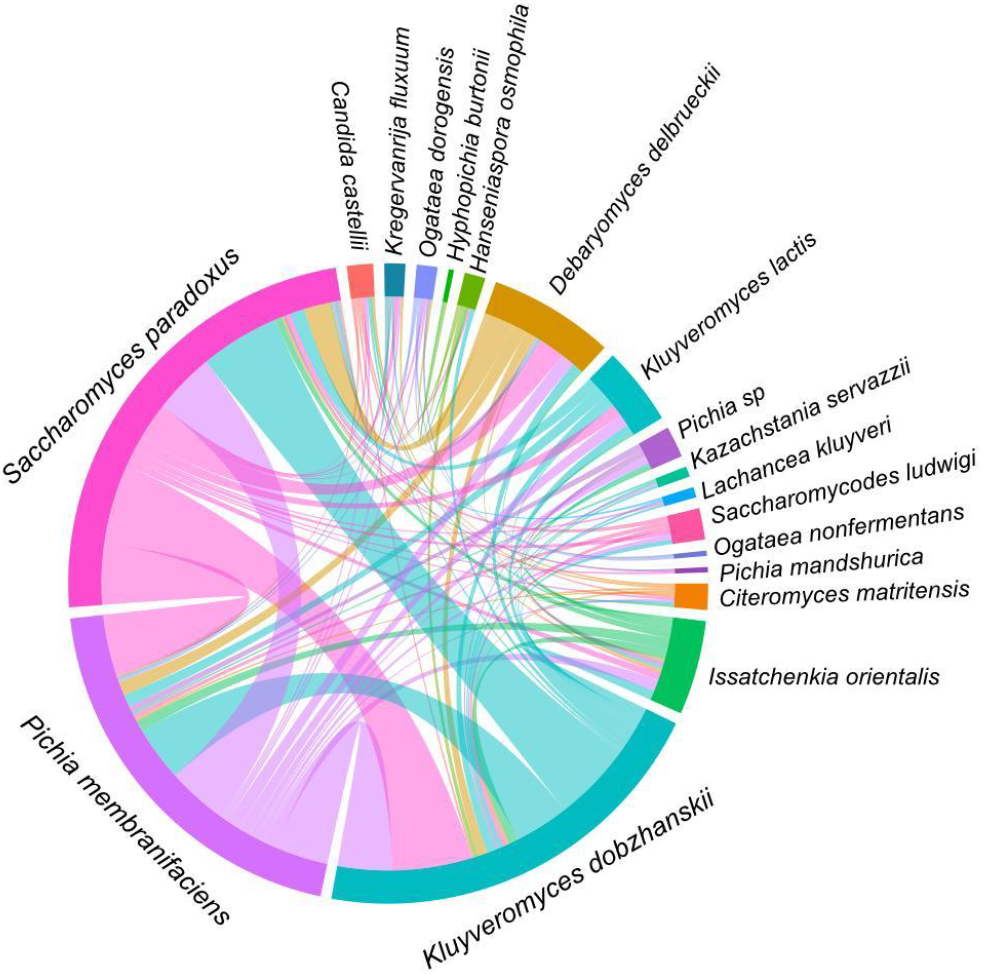
Chord diagram depicting yeast species co-occurrence, based on shared presence across sampled trees. The connections (arcs) between species indicate how frequently they co-occur, with thicker arcs representing more frequent co-occurrences.

### 2. Spatial patterns: Yeast genus dominance is correlated to environmental gradients

Next, we tested for spatial patterns in the abundance of the three most dominant yeast genera along latitudinal and longitudinal gradients. We found that the abundance of ASVs significantly decreased with longitude (going from west to east) in the genus *Kluyveromyces* (linear model: R^2^ = 0.19, p = 0.024), but not in *Saccharomyces* (R^2^ = 0.03, p = 0.370) or *Pichia* (R^2^ = 0.10, p = 0.138; **Figure 5B**). ASV abundance in *Pichia* significantly increased with latitude (going from south to north; linear model: R^2^ = 0.40, p = 0.001), but not in *Saccharomyces* (R^2^ = 0.18, p = 0.519) or *Kluyveromyces* (R^2^ = 0.04, p = 0.309; **Figure 5C**).

**Figure 5.**
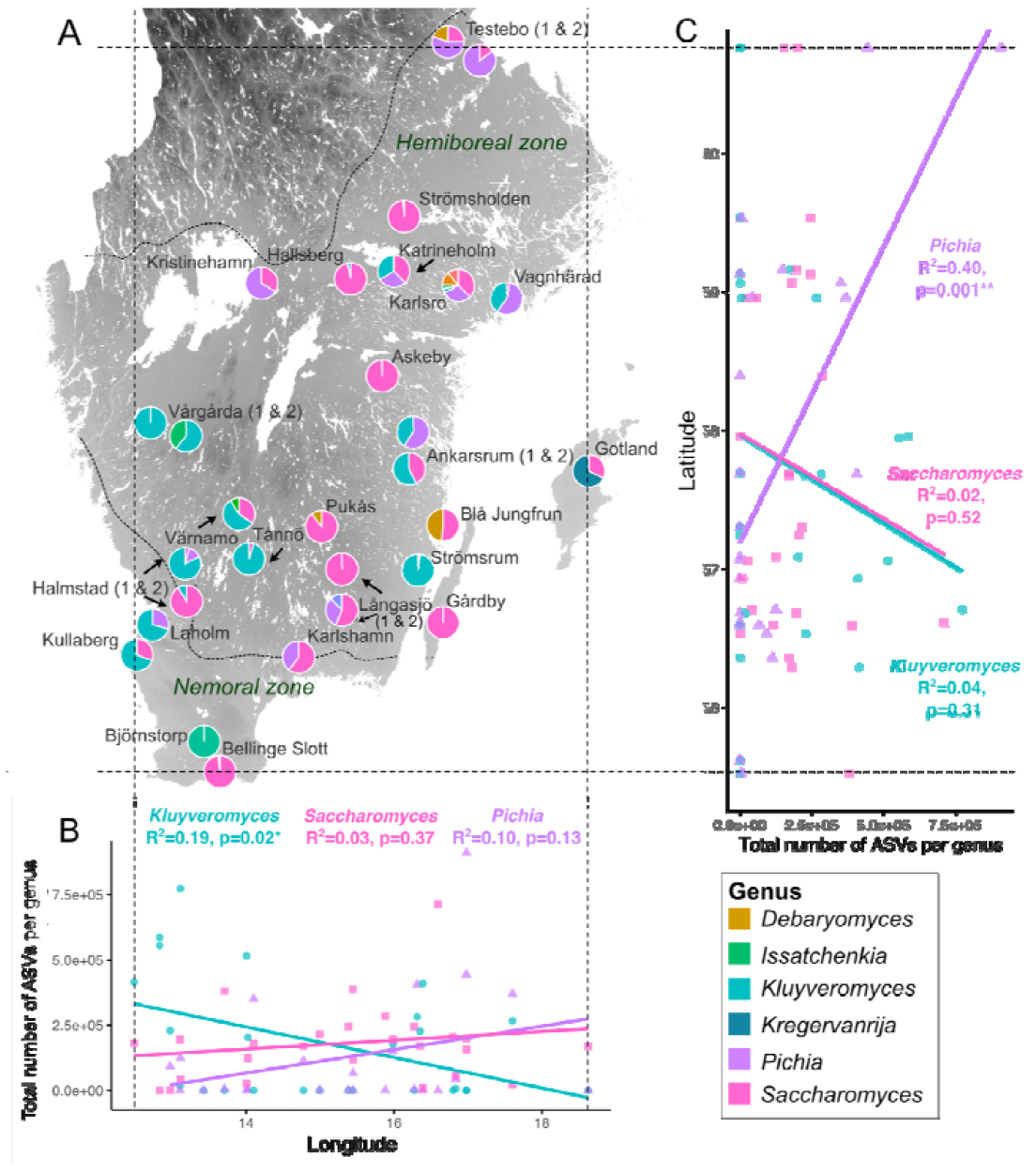
**A)** Map of sampling locations with pie charts showing the proportion of ASVs found per yeast genus. For display purposes, only genera with >2% of ASVs are shown in the legend. Dashed lines indicate the borders of hemiboreal and nemoral zones, in which oak trees were sampled. The upper line follows the northern latitudinal distribution limit of oak. **B)** Longitude, and **C)** Latitude correlated with the ASV abundance of the three most common genera isolated across all locations (*Kluyveromyces, Saccharomyces*, and *Pichia*). R-squared and p-values are indicated for each linear model per genus (p < 0.05*, <0.01**).

We then used a linear model to test whether variation in ASV abundance in *Saccharomyces, Kluyveromyces*, and *Pichia*, i.e. ‘genus’, was explained by ‘longitude’, ‘latitude’ and their interactions (minus the interactions Long:Lat and Genus:Long:Lat), which was significant (F_8,67_ = 2.99, p = 0.006). Of the individual coefficients, ASV abundance was significantly affected by longitude overall (t = - 2.45, p = 0.017) with significant differences between *Pichia* and *Kluyveromyces* (t = -2.66, p = 0.010). Interactions between *Pichia* vs. *Kluyveromyces* with latitude (t = 2.03, p = 0.047) and *Saccharomyces* vs. *Kluyveromyces* with longitude were also found to be significant (t = 2.57, p = 0.013).

Overall, these results suggest that species of *Kluyveromyces* are significantly more abundant in the south-west, that *Pichia* species are most common in the north-east, and that *Saccharomyces* species were most common in the centre of Sweden **(Figure 5)**. Despite the frequent co-occurrence of *S. paradoxus, K. dobzhanskii*, and *P. membranifaciens* (**Figure 4**), this analysis highlights the respective dominance of different genera over others along geospatial gradients.

As latitude increases going from south to north in our sampling area, minimum temperature decreases (r = -0.95, df = 71, p < 2.2e-16) and maximum precipitation increases (r = 0.64, df = 71, p = 9.05e-10; **Figure 6**). As longitude increases (going from west to east), maximum temperature (r = 0.86, df = 71, p < 2.2e-16) and minimum precipitation decrease (r = -0.73, df = 71, p = 1.74e-13). We therefore tested whether these climate metrics (maximum/minimum temperature and precipitation) also predicted variation in yeast diversity (including ASV richness, Shannon, Evenness, and the raw number of yeast species) and visualised the strength and direction of these correlations using a heatmap (**Figure 6**). We found that lower minimum precipitation negatively predicted ASV richness, suggesting that diversity is larger at drier sampling sites (r = -0.24, df = 71, p = 0.04, **Figure 6**, panel A). Note here, that this pattern may be largely caused by the high ASV richness observed for Saccharomyces (**Figure S5**).

**Figure 6.**
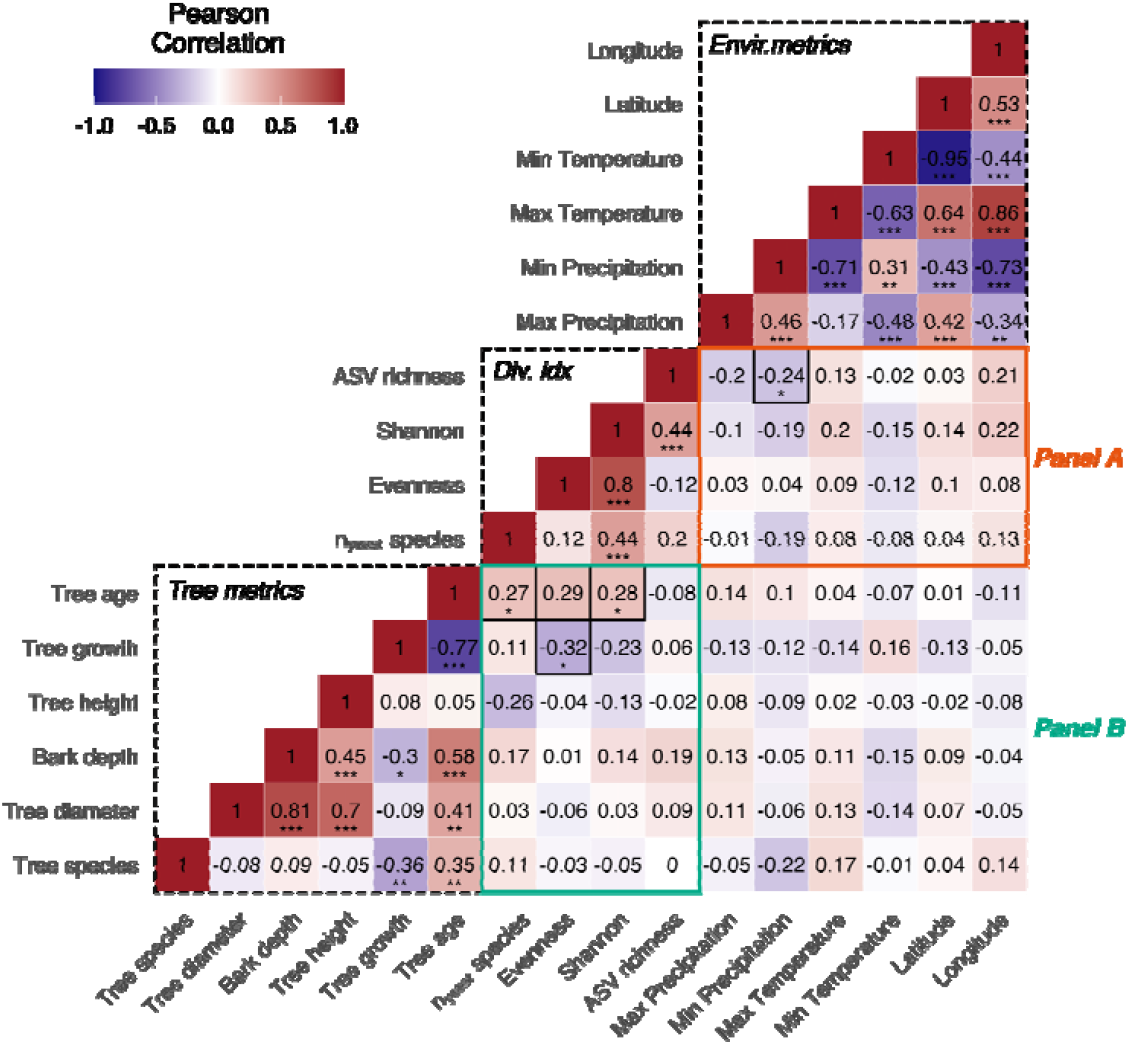
Correlation patterns between environmental variables, tree metrics and yeast diversity indices. Significance levels of Pearson correlations are indicated by asterisks (p < 0.05*, 0.01**, and 0.001***).

Analysis of the impact of temperature and precipitation on ASV abundance at the ‘genus’ level showed that lower annual minimum temperatures predicted higher ASV abundance in *Pichia* species (linear model: R = 0.26, p = 0.012; **Figure 7B**), while lower maximum temperature predicted an increase in *Kluyveromyces* ASV abundance nearly significantly (R = 0.16, p = 0.050; **Figure 7A**). Saccharomyces ASV abundance did not depend on either temperature metric. Higher minimum precipitation predicted higher ASV abundance in *Kluyveromyces* (R = 0.18, p = 0.037; **Figure 7D**), while lower maximum precipitation predicted *Saccharomyces* ASV abundance (R = 0.17, p = 0.046, **Figure 7C**). Together, these patterns suggest that the colder temperatures and higher precipitation levels in the north-eastern range limit of oak trees are well-suited for Pichia species, while the lower maximum temperatures and higher precipitation levels on the Swedish west coast are better suited for *Kluyveromyces* species. The absence of temperature effects on the distribution of *Saccharomyces* suggests that *S. paradoxus* is more thermo-generalist than the species in the other two genera.

**Figure 7.**
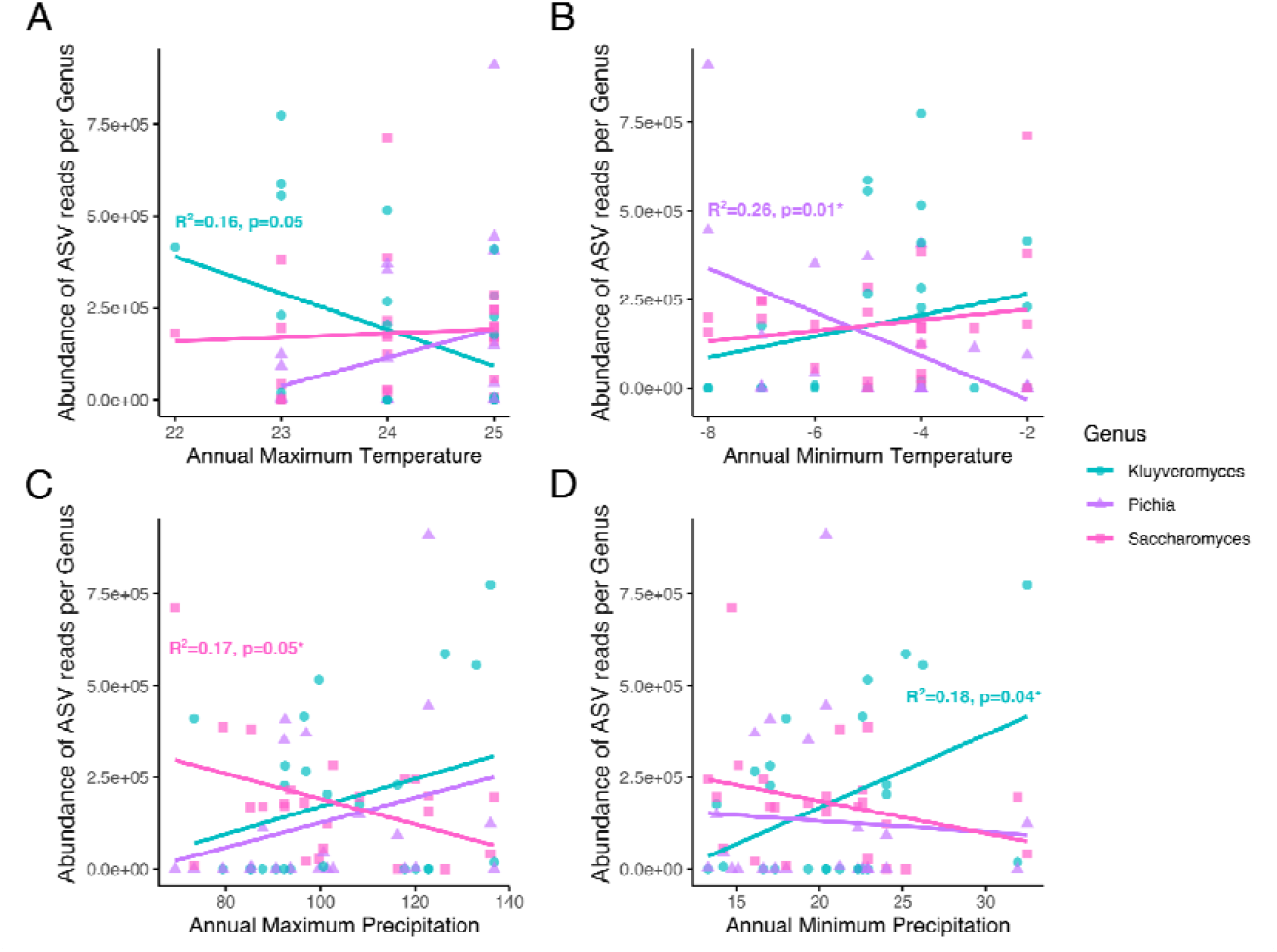
Annual **A)** maximum temperature, **B)** minimum temperature, **C)** maximum precipitation and **D)** minimum precipitation correlated with the ASV abundance (total number of ASV reads) of the three most common genera isolated across all locations (*Kluyveromyces, Saccharomyces*, and *Pichia*). R-squared and p-values are indicated for each linear model per genus if significant (or approaching significance).

### 3. Ecological drivers of yeast diversity: Older trees have richer and more balanced microbial communities

Third, we wanted to know if any ecological aspects of the oak tree hosts predicted yeast diversity. We tested for correlations between yeast diversity measures (including ASV richness, Shannon, Evenness, and the raw number of yeast species per bark sample) and several oak tree variables (tree growth, age, height, diameter, bark depth, and tree species (either *Q. robur* or *Q. petraea*)). Tree metrics are gathered in the bottom left corner of the heatmap (**Figure 6**). As expected, tree diameter was positively correlated with bark depth (r = 0.81, df = 57, p = 6.98e-15) and tree age (r = 0.41, df = 56, p =0.001). A strong negative correlation was also found between tree age and tree growth (r = – 0.77, df = 56, p = 9.17e-12), which indicates that older trees grow more slowly.

Species diversity (Shannon index, r = 0.28, df = 52, p = 0.042) and species richness (number of yeast species: r = 0.27, df = 52, p = 0.045) increased with tree age (**Figure 6**, panel B). We also found that older, more slowly growing trees have a more balanced community composition, i.e. species in the microbial communities on older trees have more evenly distributed ASV abundances (correlation between tree growth and evenness: r = -0.32, df = 45, p = 0.03).

Although only 9 trees out of the 81 sampled were identified as *Q. petraea*, they show an interesting difference in yeast diversity, although not significantly (F_1,63_ = 1.56, p = 0.156), driven by one particular tree on the island of Gotland. Of the three trees sampled on Gotland, two were *Q. robur* and one was *Q. petraea*. The latter shows a very high abundance of *Kregervanrija fluxuum*, which, to our best knowledge, is the only reported occurrence so far of this species in Sweden (**Figure 8A**). The yeast diversity on islands off Sweden’s east coast (Gotland, Gårdby, and Blå Jungfrun) was significantly different from the diversity found on the mainland (F_1,76_ = 2.27, p = 0.030). We found a strong enrichment on these islands for *Kregervanrija*, but also *Debaryomyces delbrueckii*, which was found to be highly abundant on a single tree sampled on the island of Blå Jungfrun **(Figure 8B)**.

**Figure 8.**
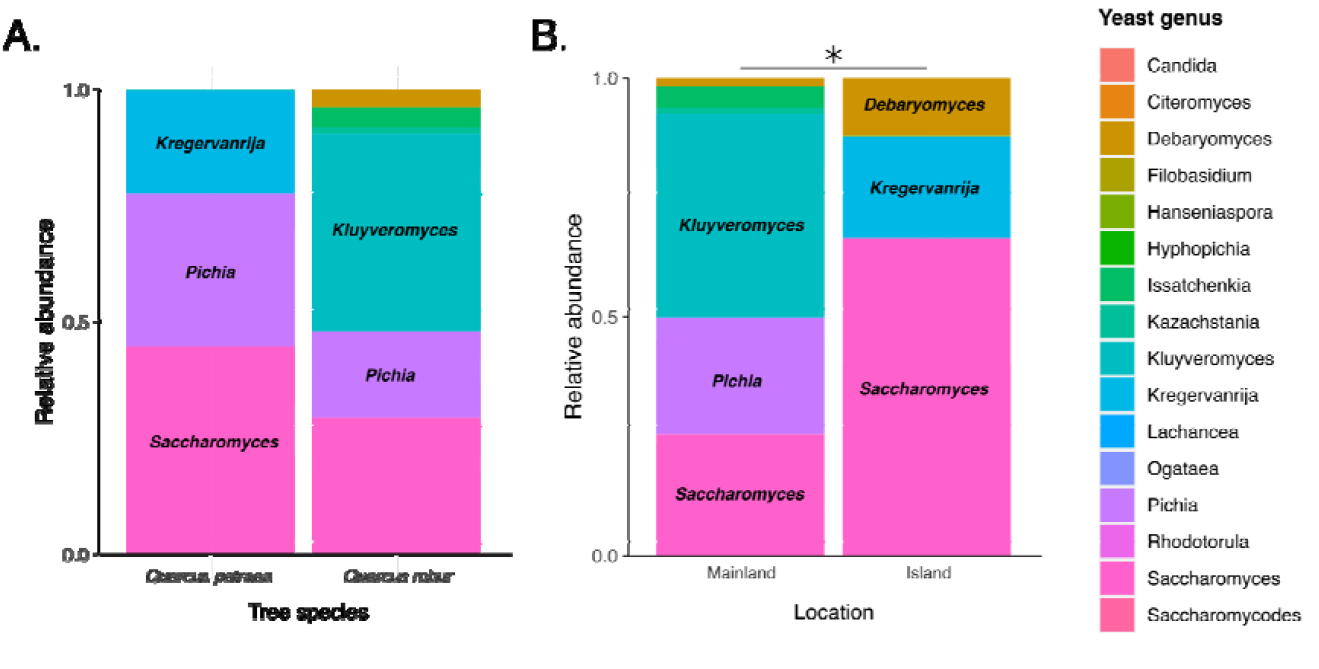
Relative abundance of yeast genera depending on **A)** host tree species and **B)** insularity. Significant differences in community composition depending on x-axis are indicated by asterisks (p < 0.05*).

### 4. The impact of temperature on the growth performance of wild yeast isolates

Finally, we tested whether the growth (maximum biomass) of the yeast strains we isolated across the sampling area was affected by temperature, using laboratory growth assays at three temperatures: 5°C, 16°C, and 35°C. We found that culturing temperature significantly affected the growth of *Kluyveromyces* collected from different longitudes and latitudes. *Kluyveromyces* strains from western Sweden grew better than strains from the east at 5°C (R^2^ =0.132, p = 0.002; **Figure 9A**), while strains collected in southern Sweden grew better than strains from the north at 35°C (R^2^ = 0.114, p = 0.005; **Figure 9B**). For *Saccharomyces*, southern strains grew better than strains from the north at 16°C (R^2^ = 0.96, p = 0.002; **Figure 9B**) but no significant effects of latitude and longitude were found when growing Saccharomyces at extreme cold or hot temperatures.

**Figure 9.**
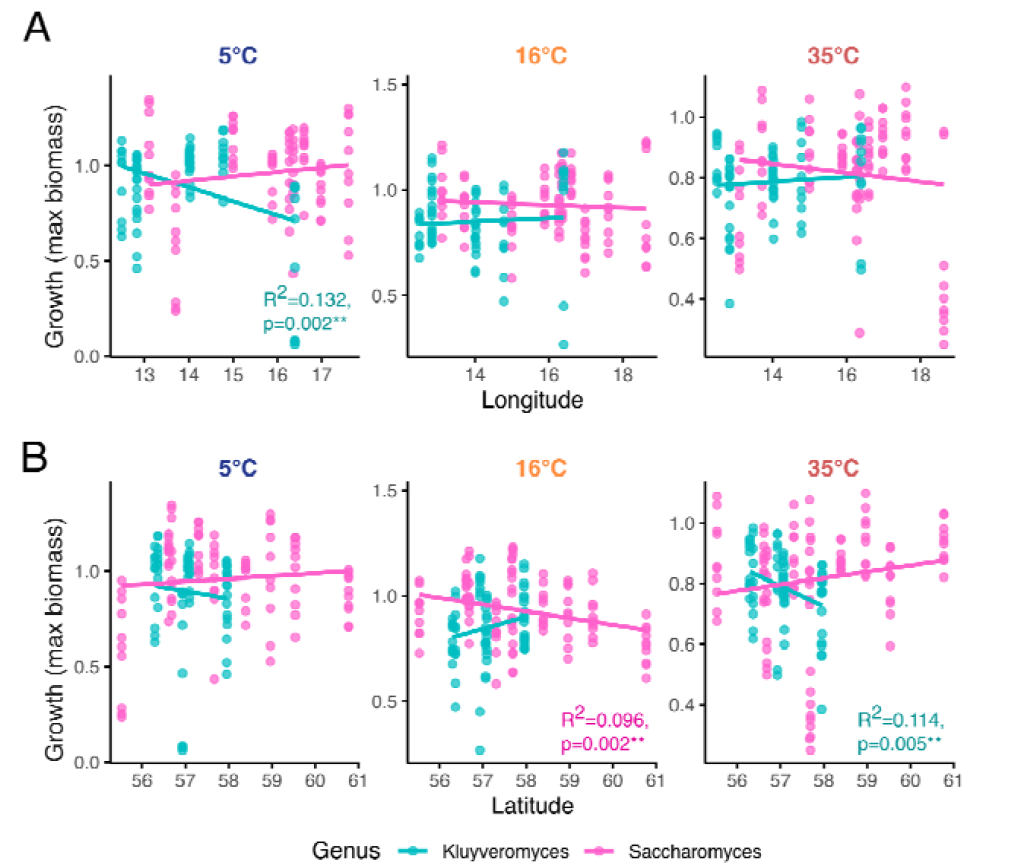
Growth (maximum biomass) of yeast strains at 5°C, 16°C and 35°C. Strains were identified by Sanger sequencing to belong to *Kluyveromyces* and *Saccharomyces*, and plotted according to their sampling location’s **A)** longitude and **B)** latitude. Each data point represents one of 10 technical replicate optical density measurements per strain and location. R-squared and p-values are indicated for each linear model per genus if significant (p < 0.01**).

## Discussion

Temperate forests are a widespread global biome and a diversity hotspot for yeast (Sniegowski et al., 2002; Sampaio & Gonçalves, 2008; Robinson et al., 2016; Mozzachiodi et al., 2022; David et al., 2024). Fermentative yeasts play crucial roles in these ecosystems. They break down organic matter and contribute to nutrient recycling by converting sugars into ethanol, carbon dioxide, and other metabolites such as esters and glycerol (Dashko et al., 2014). Despite their importance for nature and society, our knowledge of the distribution and diversity of fermentative yeast across geographically and ecologically diverse, natural environments is still scarce. Previous work suggests that oak trees are a frequent habitat for yeast (Sniegowski et al., 2002; Sampaio & Gonçalves, 2008; Robinson et al., 2016; Mozzachiodi et al., 2022) but the diversity and abundance of yeast in the northern range limit of their oak hosts has not been investigated. The temperate forests of Scandinavia are marked by a climate with strong seasonal variation between long, cold winters and mild summers. So far, most work in Scandinavia has focused on fungal communities on spruce (*Picea*) (Müller & Hallaksela, 2000) and beech (*Fagus*) (Kubart et al., 2016; Asplund et al., 2019), but no previous study has investigated whether oak trees in this region carry similar yeast communities as trees further south in the *Quercus* range. Using metabarcoding, we describe the species diversity and community composition of fermentative yeasts isolated from oak trees across a large area (covering 221,000 km^2^) in the temperate climate zone of Sweden, and test whether climatic and ecological variables predict their distribution.

### The diversity and community composition of oak-associated yeasts in Sweden

We found three main clusters of oak-associated yeast communities across southern Sweden, each dominated by species of either *Saccharomyces, Kluyveromyces* or *Pichia* (**Figure 3B**). Interestingly, microbial communities across trees were more similar when sharing a common dominant genus. Generally, the dominance of one genus did not exclude a species from another genus to co-occur in the same location, i.e. there was no strong pattern of competitive exclusion. However, in some locations, only a single genus was detected (small dots in **Figure 3B**).

One species pair co-occurred more frequently than expected by chance: *Debaryomyces delbrueckii* and *Saccharomyces paradoxus* (Figure S4), which suggests they have compatible ecological requirements or enter beneficial metabolic interactions. A study from temperate woodlands in North America has found S. paradoxus to often co-occur with *S. cerevisiae* (Sweeney et al., 2004). Although *S. cerevisiae* is a cosmopolitan species due to its long history of domestication for alcoholic beverage production, we did not detect *S. cerevisiae* in Sweden. However, just as *S. cerevisiae, D. delbrueckii* (also called Torulaspora delbrueckii) has been domesticated for winemaking (Albertin et al., 2014) and has high tolerance to extremely low temperatures and freezing (Alves-Araújo et al., 2004). Thus, *D. delbrueckii* may occupy a similar ecological niche in Sweden as *S. cerevisiae* elsewhere in the world, but may have adaptive advantages at higher latitudes due to its extreme cold tolerance. Other species pairs in our collection rarely or never overlapped in the same location (e.g. *D. delbrueckii* with *K. dobzhanskii* or S. paradoxus with Pichia sp.), perhaps due to competition or distinct habitat preferences. The positive co-occurence of *D. delbrueckii* with *S*.*paradoxus*, and its negative co-occurence with *K. dobzhanskii*, reinforce our hypothesis that Saccharomyces and Kluyveromyces have different niches in Sweden (**Figure 5**).

We also identified non-conventional yeast such as Saccharomycodes, used in the production of alcohol-free beer (Montanari et al., 2009), Kazachstania, a yeast with potential use in biorefineries (Balarezo-Cisneros et al., 2023), *Isaatchenkia*, a halophilic yeast with the capacity to grow at low-pH conditions (Matsushika et al., 2016), and *Kregervanrija*, a yeast that weakly ferments sugars (Kurtzman, 2011). Some of the species we detected are also found in human-associated environments, e.g. during the spontaneous fermentation of grape juice. Non-*Saccharomyces* yeasts such as *Hanseniospora, Candida, Pichia*, and *Kluyveromyces* are predominant during the initial phase of alcoholic fermentation, before the conversion of sugars into ethanol by *Saccharomyces* (Jolly et al., 2014). For instance, *P. membranifaciens* is commonly used in industrial applications because it produces “killer toxins” that help control the growth of spoilage yeast and filamentous fungi (Santos & Marquina, 2004).

Since the enrichment medium we used for strain isolation prior to metabarcoding favours the growth of Saccharomycodaceae yeasts, and ITS sequencing protocols can add additional bias (De Filippis et al., 2017), we cannot rule out the possibility that other yeast species are also present on Swedish oak. Our collection of oak-associated fermentative yeast from southern Sweden, including 1150 cryo-preserved strains across all sampling locations, is available upon request for future experimental work in ecology and evolution, or for potential uses in industry.

### The abundance and distribution of oak-associated yeasts in Sweden

Our results reveal interesting patterns in the abundance and distribution of oak-associated yeast across a longitudinal gradient in southern Sweden, and suggest that climate variation from west to east and south to north, especially in temperature and precipitation (**Figure 7**), contribute to this. Both coasts experience an oceanic climate, with more extreme seasonal temperature fluctuations in the east than on the milder west coast (Köppen & Geiger, 1930). The west coast in particular is characterised by a more humid climate. Going northwards, the climate gets colder and wetter, and the vegetation changes from agricultural lands to deciduous forests, mixed forests, and even some taiga. Our findings suggest genus-specific responses to these climatic gradients (**Figure 5B**). *Kluyveromyces* showed a strong negative correlation with longitude with higher abundance in western regions, while Pichia dominated in the east, and *Saccharomyces* was more common in central areas. Compared to *Pichia, Kluyveromyces* may be less tolerant to the larger temperature swings on the east coast. These differences in the spatial distribution of yeast genera across Sweden, together with their overlap in some locations (**Figure 3B**), suggest that niche-defining ecological and climate factors are more likely to shape microbial community composition than competitive exclusion. In addition to climate, factors such as soil composition and competition from other microbes, which we have not assessed here, also likely influence the observed distribution patterns (Brockett et al., 2012; Kowallik et al., 2015; Mundra et al., 2021).

### Effects of temperature on the growth of yeast isolates from the wild

Previous studies have shown that species of both *Kluyveromyces* and *Pichia* have a wide range of critical thermal minima and maxima, from 5°C to 45°C (Dickson et al., 1979; Slininger et al., 1990; Nambu-Nishida et al., 2017) and their frequent isolation from plants and insects in North America and Russia (Sukhotina et al., 2006) suggests that they are resilient to a large range of temperatures. Our assays revealed significant effects of temperature on the growth of fermentative yeast collected from different longitudes and latitudes in Sweden, especially in the genus *Kluyveromyces* (**Figure 9**). The better cold performance (at 5°C) of *Kluyveromyces* strains collected in the west compared to strains from the east, suggests they are able to perform well in the relatively milder winter climate on the west coast, while the better performance of southern vs. northern strains at hot conditions (35°C) indicates that southern strains are well adapted to the warm summer temperatures typical for this region, including occasional heat waves. This pattern is consistent with our finding that the more moderate climate in the southwest predicts a significantly higher abundance of *<u>Kluyveromyces</u>* (**Figure 5**). In agreement with this, a study in North America showed that *K. dobzhanskii* was most frequently isolated at moderate temperatures (10 - 20°C) but less often at 30°C (Sylvester et al., 2015).

In *Saccharomyces*, the absence of significant effects of longitude and latitude on growth at extreme temperatures (**Figure 9**) is in line with the previously reported thermogeneralist performance of *S. paradoxus* (Sniegowski et al., 2002; Robinson et al., 2016) - the most abundant yeast species in our sampling area (**Table 1, Figure 3**). *S. paradoxus* is, after *S. cerevisiae*, the most cosmopolitan species of *Saccharomyces* and is frequently isolated from trees, soil, and bark of deciduous trees in the Northern Hemisphere (Charron et al., 2014). The ability of *Saccharomyces* to grow both at warm and cold temperatures, for instance through the production and subsequent use of glycerol as a cryo-preservative (Hohmann, 2002; Koh, 2013), may give them a selective advantage over other fermentative yeasts in the generally cold climate of Sweden (Sweeney et al., 2004; Salvadó Z. et al., 2011).

### Conclusions

Our study provides insights into the diversity and distribution of fermentative yeasts in the northern range limit of their oak tree hosts. To our knowledge, our data provide the first evidence that i) older oak trees supported greater yeast species richness, and ii) that the relative strain frequencies in each tree-specific microbial community are more balanced, the older the tree is. This suggests that tree age plays an important role in establishing more stable microbial communities, allowing multiple species to coexist in similar abundance. We found that communities were largely dominated by species of the Saccharomycetaceae family, with notable longitudinal and latitudinal gradients in their composition, which are associated with climate variables, especially temperature and precipitation. Future research on the underlying genetic basis of thermal tolerance of these wild yeast isolates may shed light on their vulnerability to future climate change. This is important as changes in microbial community compositions can have cascading effects through the food web, and potentially affect the diversity of entire ecosystems (Pörtner & Farrell, 2008) and their services (Cavicchioli et al., 2019). Together, our findings contribute to a better understanding of the diversity of natural yeast communities and the drivers of their distribution in the environment.

## Acknowledgements

We thank the oak bark collection team, including Genoveva Elisabeth Zimmermann, Jonas Lundqvist, and Sofia Rouot for her help with processing samples in the lab. The work was funded by the Swedish Research Council (grant 2022-03427) and the Knut and Alice Wallenberg Foundation (2017.0163) to RS. The authors acknowledge support from the National Genomics Infrastructure in Stockholm funded by the Science for Life Laboratory, and the SNIC/Uppsala Multidisciplinary Center for Advanced Computational Science for assistance with sequencing and access to the UPPMAX computational infrastructure. Computation and data handling were enabled by resources in project NAISS 2023/22-116 provided by the National Academic Infrastructure for Supercomputing in Sweden (NAISS) at UPPMAX, funded by the Swedish Research Council through grant agreement no. 2022-06725.

