## Supplement Table S1, Supplement Figure S1 to S5 for "Fermentative yeast diversity at the northern range limit of their oak tree hosts"

### Supplementary Material

**Table S1.** Primer sequences for ITS-region PCR

| Name | Forward sequence (5′-3′) | Reverse sequence (5′-3′) |
|---|---|---|
| NGI overhangs - ITS1 (PCR1) | ACACTCTTTCCCTACACGACGCTCTTCCGATCT – TCCGTAGGTGAACCTGCGG | GTGACTGGAGTTCAGACGTGTGCTCTTCCGATCT – GCTGCGTTCTTCATCGATGC |

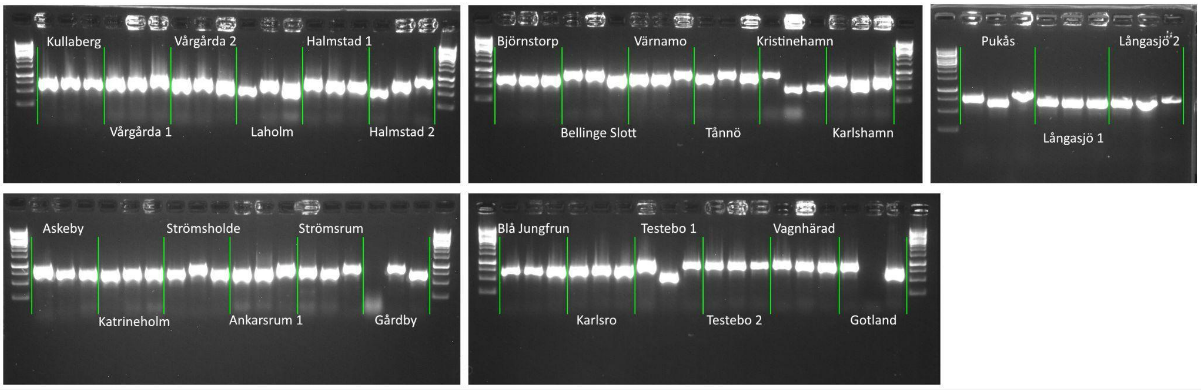


**Figure S1.** Confirmation of amplification of the ITS region via PCR. Sampling locations are organized along longitude (from west to east).

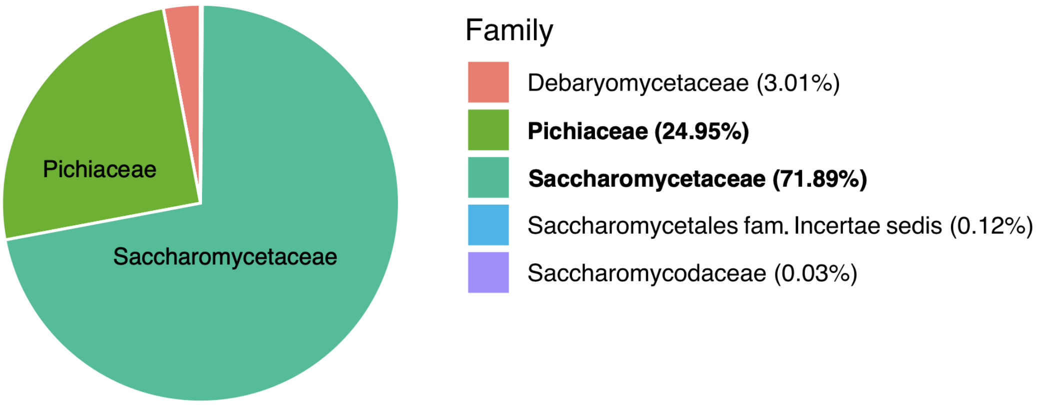


**Figure S2.** Percentage of ASVs mapping to each family across all samples.

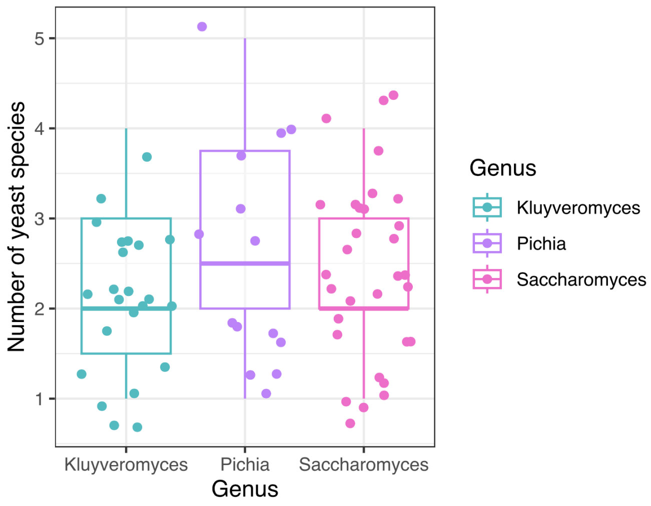


**Figure S3.** Number of species in the three clusters that group the dominant genera. Boxplot extremities represent minimum and maximum values, whereas the box itself is composed of the first quartile, median (thick line), and third quartile. A jitter effect was added for better visibility of data points.

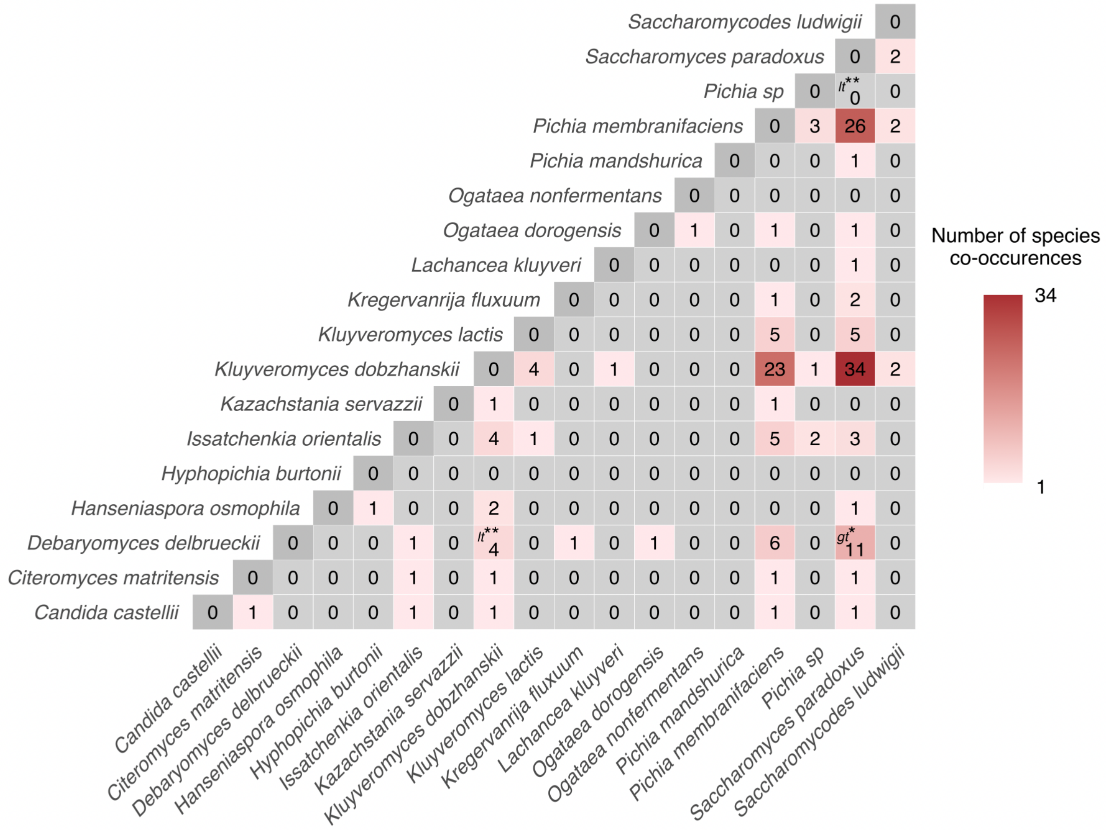


**Figure S4.** Co-occurrence heatmap between yeast species detected across all trees and sites based on probabilistic assessment of the observed *vs* expected co-occurrence frequencies. Pairs of species with significantly fewer (p-values 'lesser than', *lt*) or more co-occurrences (p-value 'greater than', *gt*) than expected by chance are indicated by asterisks (p < 0.05* and 0.01**).

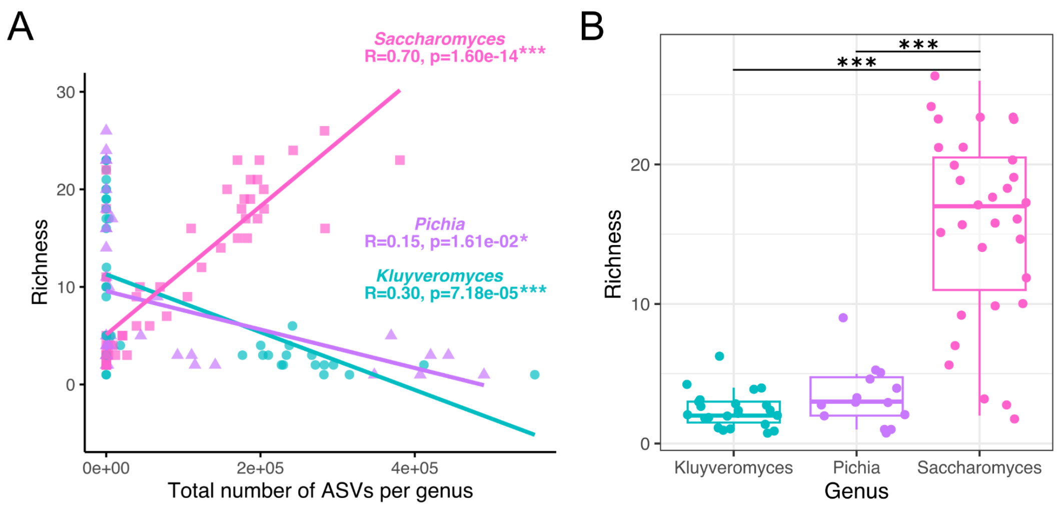


**Figure S5.** ASV richness of species in the three dominant genera, **A)** correlated with ASV abundance and **B)** represented as boxplots. R-squared and significance level of associated p-values for each linear model per genus are indicated by asterisks (p < 0.05*, 0.01**, and 0.001***) on A, while on B Asterisks indicate statistical significance using Kruskal-Wallis followed by pairwise Wilcoxon post hoc tests with Bonferroni corrections (adjusted p < 0.05*, 0.01**, and 0.001***).
